# Synapse development is regulated by microglial THIK-1 K^+^ channels

**DOI:** 10.1101/2021.04.05.438436

**Authors:** Pablo Izquierdo, Hiroko Shiina, Chanawee Hirunpattarasilp, Huma Sethi, David Attwell

## Abstract

Microglia are the resident immune cells of the central nervous system. They constantly survey the brain parenchyma for redundant synapses, debris or dying cells, which they remove through phagocytosis. Microglial ramification, motility and cytokine release are regulated by tonically active THIK-1 K^+^ channels on the microglial plasma membrane. Here, we examined whether these channels play a role in phagocytosis. Using pharmacological blockers and THIK-1 knockout (KO) mice, we found that lack of THIK-1 activity reduced microglial phagocytosis, which may result in impaired pruning of synapses. In hippocampus, mice lacking THIK-1 expression had an increased number of glutamatergic synapses during development. This resulted from an increased number of presynaptic terminals, due to impaired removal by THIK-1 KO microglia. In microglia in brain slices from fresh human biopsies, modulating THIK-1 function had effects similar to those in rodents: blocking THIK-1 rapidly reduced microglial process ramification and increased synaptic density. The dependence of synapse number on THIK-1 K^+^ channels, which control microglial surveillance and phagocytic ability, implies that changes in THIK-1 expression level over the lifespan or in disease states may contribute to altering neural circuit function.

**Significance:** Microglia are the brain’s resident immune cells, surveying it with motile processes, which can remove pathogens but also prune unnecessary junctions between the neurons (synapses). A potassium channel, THIK-1, in the microglial membrane allows efflux of potassium from these cells, and thereby regulates their membrane voltage as well as their process motility and release of inflammatory mediators. Here, using THIK-1-blocking drugs and THIK-1-deficient mice, we demonstrate that THIK-1 controls removal of synaptic material by microglia, which reduces the number of functional synapses. We also show that blocking THIK-1, as some anaesthetics do, affects microglial structure and increases the number of synapses in living brain slices from both rodents and humans, and could thus alter network function in the brain.

## INTRODUCTION

Microglia are the primary immune cells in the brain parenchyma, accounting for 5-12% of all cells, with a particularly high density in the hippocampus (1). Their functions are controlled by a plethora of ion channels and receptors on their plasma membrane (2). Even so-called ‘resting’ microglia are constantly scanning the surrounding brain tissue, by extending and retracting their processes. Baseline surveillance is regulated independently of targeted chemotaxis, and is controlled by membrane voltage, the resting value of which (around −40 mV) is set by the two-pore domain halothane-inhibited K^+^ (THIK-1) channel, the main K^+^ channel expressed by microglia *in situ* (3).

Microglia can internalize and degrade targets that they have previously detected and contacted. This process, referred to as phagocytosis, occurs throughout life in both physiological and pathological conditions. Microglia can phagocytose both dead and viable cells (4), myelin sheaths (5) and amyloid debris (6).

However, microglia are more than mere cleaners of the brain parenchyma. These cells are key for normal brain development (7) and their processes interact with various neuronal compartments. While microglial contacts at non-synaptic regions regulate dendrite branching and axonal pathfinding during development as well as neuronal excitability, contacts at synaptic sites control synapse formation, strength, plasticity and elimination (8–10). Microglia-synapse contacts can lead to formation of dendritic spines as well as new filopodia in spine heads (11, 12). On the other hand, microglia-mediated circuit pruning also occurs as redundant or less active synapses are removed (13, 14). Synapse phagocytosis is controlled by a fine balance between “eat-me” and “don’t-eat-me” signals, which respectively promote removal (e.g., C1q or C3/CD11b (15)) or protect synapses from pruning (e.g., CD47 (16)). A number of signalling pathways are involved in microglia-mediated removal of synapses during development, including complement factors in the visual system (14, 17) or the fractalkine receptor (13, 18) and TREM2 (19) in the hippocampus.

Microglial THIK-1 K^+^ channels maintain ramification of microglial processes and their surveillance of the brain (3), so they could promote the elimination of synapses during development by phagocytosis (or trogocytosis (12)). Immune cells have been shown to hyperpolarise when phagocytosing targets (20) so, in addition to promoting microglial-synapse interactions, the hyperpolarization maintained by microglial THIK-1 channels might also promote phagocytosis directly. Regulation of phagocytosis by membrane potential could also explain why phagocytosis is impaired in TREM2 knockout mice where microglia are depolarized (21). Because developmental mechanisms are thought to be reactivated during ageing and disease (22–26), understanding the mechanisms controlling microglial phagocytosis may have therapeutic value in the mature brain.

Here, we identify membrane K^+^ flux as a factor regulating microglial phagocytosis and show that block or deletion of THIK-1 channels reduces microglial phagocytosis *in situ*. THIK-1 deletion results in decreased microglial uptake of presynaptic material in the developing hippocampus, and thus an increased number of functional glutamatergic synapses. Potentially importantly for clinical translation, block of THIK-1 channels in human microglia recapitulated its effects in rodents, reducing microglial ramification and increasing synapse numbers.

## MATERIALS AND METHODS

### Rodent procedures

All animal procedures were performed in accordance with the UK Animals (Scientific Procedures) Act 1986 (Home Office License 70/8976). Rodents were maintained on a 12-hour/12-hour light/dark cycle, and food and water were available *ad libitum*. Mice (postnatal day P17–19, and P26–32) were housed in individually ventilated cages and Sprague-Dawley rats (P12–13) were kept in open-shelf units. Animals of both sexes were sacrificed by cervical dislocation followed by decapitation, or by an overdose of pentobarbital sodium (Euthatal, 200 µg/g body weight) injected intraperitoneally before transcardial perfusion-fixation of tissue with 4% paraformaldehyde (PFA).

For mouse experiments, THIK-1 knockouts (Kcnk13-IN1-EM1-B6N) were generated by MRC Harwell as previously described in detail (3). Briefly, a single nucleotide insertion in the gene encoding the THIK-1 channel protein (*Kcnk13*) leads to a premature stop codon, producing a truncated protein that fails to form a channel.

### Acute brain slicing

For microbead assays, 250 µm hippocampal slices were prepared on a Leica VT1200S vibratome in ice-chilled slicing solution containing (in mM): 124 NaCl, 2.5 KCl, 26 NaHCO_3_, 1 NaH_2_PO_4_, 10 glucose, 1 CaCl_2_, 2 MgCl_2_ and 1 kynurenic acid. Osmolarity was adjusted to ∼295 mOsM and pH set to 7.4 when bubbled with 5% CO_2_.

For electrophysiology and human tissue experiments, 300 µm hippocampal slices from mice and cortical slices from human samples were prepared in ice-cold N-methyl-D-glucamine (NMDG)-based slicing solution (to reduce cell swelling), containing (in mM): 93 NMDG, 2.5 KCl, 20 HEPES, 30 NaHCO_3_, 1.2 NaH_2_PO_4_, 25 glucose, 0.5 CaCl_2_, 10 MgCl_2_, 5 sodium ascorbate, 2.4 sodium pyruvate and 1 kynurenic acid. The osmolarity was adjusted to ∼300 mOsM, and pH set to 7.4. Slices were immediately transferred to warmed (35°C) slicing solution for 20 min, and then to storage solution at room temperature containing (in mM): 92 NaCl, 2.5 KCl, 20 HEPES, 30 NaHCO_3_, 1.2 NaH_2_PO_4_, 25 glucose, 2 CaCl_2_, 1 MgCl_2_, 5 sodium ascorbate, 2.4 sodium pyruvate and 1 kynurenic acid. The osmolarity was adjusted to ∼300 mOsm, and pH set to 7.4.

### Slice incubation experiments

Brain slices from human or rat were incubated in artificial cerebrospinal fluid (aCSF) containing (in mM): 140 NaCl, 2.5 KCl, 10 HEPES, 1 NaH_2_PO_4_, 10 glucose, 2 CaCl_2_ and 1 MgCl_2_. aCSF was supplemented with 50 µM tetrapentylammonium (TPA) or 50 µM bupivacaine when needed. The osmolarity was adjusted to ∼295 mOsM and the pH set to 7.4. Slices were placed in small chambers and incubated at 35°C for 40 min prior to fixation in 4% PFA for 1 hour (human) or 45 min (rat) at room temperature and immunostaining as described below.

### Microbead phagocytosis assay

Hippocampal brain slices were allowed to recover at room temperature for 2.5 hours so that the microglia would activate and become more phagocytic (27–29). They were then transferred to 24-well plates and incubated with 3 µm serum coated, FITC-labeled resin microbeads (Sigma 72439; 1.7×10^7^ beads per well in aCSF) for 1.5 hours in a cell culture incubator at 35°C (or at 4°C, as a negative control). Bead suspensions were supplemented as indicated with the following drugs: bupivacaine (Sigma B5274), charybdotoxin (Anorspec 28244), cytochalasin D (Sigma 22144), MRS2578 (Cayman CAY19704), TPA (Sigma 258962) or tetrodotoxin citrate (TTX, Abcam ab120055). In addition, 30 min prior to applying the bead suspension (i.e., during the last 30 min of the recovery period), slices were pre-incubated with the drugs at the same concentration as that contained in the bead suspension. For high [K^+^]_o_, aCSF containing 120 mM KCl (replacing 117.5 mM NaCl with KCl) was used. Following incubation, slices were rinsed in cold phosphate-buffered saline (PBS), fixed in 4% PFA for 45 min at room temperature and immunostained as described below. Confocal stacks were obtained at 2 μm z-step intervals using a Zeiss LSM700 microscope with a Plan-Apochromat 20×/0.8 objective. To assess phagocytosis, the percentage of phagocytic microglia (i.e., Iba1^+^ cells which had internalized >1 FITC^+^ beads) was calculated. A total of 4–30 slices (3–5 fields of view averaged per slice) from 2–6 animals were used per condition. All imaging and analyses were done with the researcher blind to genotype and treatment.

### Immunohistochemistry

Free-floating 250 µm hippocampal slices were permeabilized and blocked for 2 hours at room temperature in a solution containing 10% normal horse serum and 0.02% Triton X-100 in PBS, followed by incubation with primary antibodies in blocking buffer for 12 hours at 4°C with agitation (mouse anti-Bassoon (1:300, Novus NB120-13249), rabbit anti-Homer1 (1:300, Synaptic Systems 160002), rabbit anti-Iba1 (1:500, Synaptic Systems 234003), mouse anti-vGluT1 (1:300, Abcam ab134283)). Following four 10 min washes in PBS, secondary antibodies diluted 1:1000 in blocking buffer (donkey anti-mouse Alexa 488, donkey anti-rabbit Alexa Fluor 555 or 647, Invitrogen) were applied for 4 hours at room temperature with agitation. Finally, slices were washed in PBS and mounted.

### Synapse quantification

Brain slices were imaged (102 µm x 102 µm) at 3–6 μm from the slice surface using a Zeiss LSM700 microscope with a Plan-Apochromat 63×/1.4 objective. Three confocal images from the CA1 *stratum radiatum* region (at 1.5 μm intervals) were taken per brain slice, and 5 brain slices were taken per animal. Images were analysed individually and then averaged across stacks and brain slices to obtain animal means. After background subtraction (with a 10-pixel rolling ball average), marker areas were quantified using a custom-based intensity threshold protocol with FIJI, and the Analyze Particles function was used to quantify puncta number and areas. For thresholding, sample images were first manually thresholded for each channel blinded to condition and genotype, and a suitable range above threshold was established which was then kept constant throughout (Bassoon: 15-255; Homer1: 30-255; images were 8-bit). Size exclusion (>1.2 μm^2^, <0.05 μm^2^) was applied to exclude any objects unlikely to represent synaptic puncta. Synapses were defined by the presence of overlapping presynaptic-postsynaptic puncta (30). Presynaptic colocalization with microglia was analysed as the total area of Bassoon puncta within the Iba1-stained cell area after thresholding. For human synapse analysis, images were processed by super-resolution radial fluctuation (SRRF) analysis (31) to improve resolution prior to thresholding and quantification. All imaging and analysis were done with the researcher blind to genotype or treatment.

### Cell density analysis

Brain slices were imaged (640 µm x 640 µm) using a Zeiss LSM700 microscope with a Plan-Apochromat 20×/0.8 objective. Cell density and spatial distribution were analysed as in (32). Briefly, cell counts were performed to obtain cell density as well as the average nearest-neighbour distance between cells (NND) and their regularity index. The latter is the ratio of the mean NND to the standard deviation of the NND for the whole population of cells and describes how regular is the spacing of microglia. One maximum-projected z-stack (3 µm depth) from the CA1 *stratum radiatum* region was analysed per slice, and 5 slices were averaged per animal.

### Sholl analysis

Confocal stacks (at 0.34 µm z-step intervals) of Iba1-stained microglia were obtained from the CA1 *stratum radiatum* using a Zeiss LSM700 microscope with a Plan-Apochromat 63×/1.4 objective. Three-dimensional cell reconstructions were performed using the automatic cell tracing tool ‘Neuron Tracing v2.0’ on Vaa3D (vaa3d.org). After the reconstructions were manually verified, they were analysed using custom-written MATLAB software described in (3) and available from github.com/AttwellLab/Microglia. Briefly, a series of concentric spheres were drawn at 5 µm intervals from the centre of the cell and a profile of branching points was generated across the Sholl spheres to assess cell ramification and overall process architecture. For each condition 85–90 cells were analysed. All imaging and analysis were done with the researcher blind to treatment.

### Electrophysiology

Slices were individually transferred to the recording chamber and perfused at 3-5 ml/min with aCSF, which was maintained at 32–34°C. Pyramidal neurons in the CA1 region of the dorsal hippocampus were selected visually using an Olympus 60×/0.9 water-immersion objective in combination with differential interference contrast (DIC) optics. Cells were recorded in the whole-cell voltage-clamp configuration with glass patch-pipettes (resistance in the bath solution 2–4 MΩ). Junction potentials (−10 mV) were corrected for. Recorded signals were sampled and digitized at 20 kHz, filtered at 4 kHz, and then further filtered offline at 2 kHz for analysis and data presentation. During the entire course of recording, access resistance was monitored by periodically applying a −5mV voltage pulse. Cells were excluded from analysis if the access resistance changed by more than 20% during the course of an experiment.

To study excitatory synapses, pipettes were filled with internal solution containing (in mM): 132.3 K-gluconate, 7.7 KCl, 4 NaCl, 0.5 CaCl_2_, 10 HEPES, 5 EGTA, 4 MgATP, and 0.5 Na_2_GTP (pH 7.2–7.3). The calculated reversal potential for Cl^-^ (E_Cl_) with these solutions was ∼ −62mV, and a holding potential (V_h_) of ∼ −65mV was used. Thus, at V_h_=-65mV, Cl^-^-mediated IPSCs were nearly invisible and cation-mediated EPSCs are inward. To isolate single vesicular events, 500 nM TTX was applied, and the frequency and amplitude of EPSCs in TTX were monitored. To assess whole cell glutamate receptor-mediated currents, cells were whole-cell voltage-clamped at −40mV and recorded in the presence of the GABA_A_R antagonist Gabazine (10 μM, Tocris 1262). N-methyl-D-aspartic acid (NMDA, 10 μM; Tocris 0114) or kainic acid (Tocris, 1 μM; Tocris 0222) were bath-applied sequentially (ensuring that holding current returned to the original control level before subsequent drug application, 10–15 min), and the resulting change in current was measured. Peak shifts in holding current induced by each drug were reported as the appropriate glutamate receptor-mediated current.

To study inhibitory synapses, pipettes were filled with internal solutions containing (in mM): 140 K-gluconate, 1.4 NaCl, 1.5 MgCl_2,_ 0.5 CaCl_2_, 10 HEPES, 0.2 EGTA, 4 MgATP, and 0.5 Na_2_GTP (pH 7.2 – 7.3). E_Cl_ calculated for these solutions was ∼-85mV and a holding potential (V_h_) of −50mV was used. Thus, at V_h_=-50mV, Cl^-^-mediated IPSCs are outward and cation-mediated EPSCs are inward. All drugs were dissolved in aCSF and bath applied. To isolate single vesicular events, 500 nM TTX was applied, and the frequency and amplitude of IPSCs in the TTX were monitored.

### Electrophysiology data analysis

The frequencies and amplitudes of spontaneous and miniature EPSCs and IPSCs (sEPSCs, sIPSCs mEPSCs, and mIPSCs, respectively) were measured using automatic detection of these events with Mini Analysis 6.0.7 (Synaptosoft) software followed by inspection of individual events for analysis. For assessment of synaptic current frequencies, events during the 180 s immediately before TTX application (defined as the baseline period) were compared to mean frequencies observed from 100 s after TTX application onset (over 180 s). The change of mean current induced by NMDA and kainic acid was calculated by defining the mean current in 20 s segments for control solution and at the peak of the NMDA and kainic acid applications by making histograms of all data points, and then fitting a Gaussian distribution to each histogram to define the mean current (using Clampfit 10.4, Molecular Devices).

### Human biopsy tissue

Human tissue data were obtained while investigating a possible role for microglia in regulating pericyte-mediated control of cerebral blood flow by altering the number of functional synapses. Living tissue was obtained from the National Hospital for Neurology and Neurosurgery (Queen Square, London). Cortical biopsies were taken from subjects aged 29–74 undergoing glioma resection. Healthy brain tissue overlying the tumour, which would otherwise have been discarded, was used. Ethical approval was obtained (REC number 15/NW/0568, IRAS ID 180727 (v3.0), as approved on 9-10-2018 for extension and amendment to include cells interacting with pericytes) and all patients gave informed written consent. All tissue handling and storage were in accordance with the Human Tissue Act (2004).

### Statistics

Data are presented as mean ± standard error of the mean (s.e.m.). Data normality was assessed using the D’Agostino-Pearson test. Statistical significance (taken as p<0.05) was assessed using unpaired two-tailed Student’s t-tests (Fig. 2*I*, 3*C*, 3*F-G*, 4*F*), Mann-Whitney tests (Fig. 1*E*, 2*B-G*, 3*B*, 3*D-E*, 4*C*, S3, S4, S5, S6*C*, S7*C*, S7*F*) or one-way analysis of variance (ANOVA) followed by Dunnett’s post-hoc tests for individual comparisons (Fig. 1*C*, 4*D*, S2, S6*D*, S7*D*). All statistical analysis was performed in Microsoft Excel 2016, GraphPad Prism 7 and Sigmaplot 11.

**Fig. 1.**
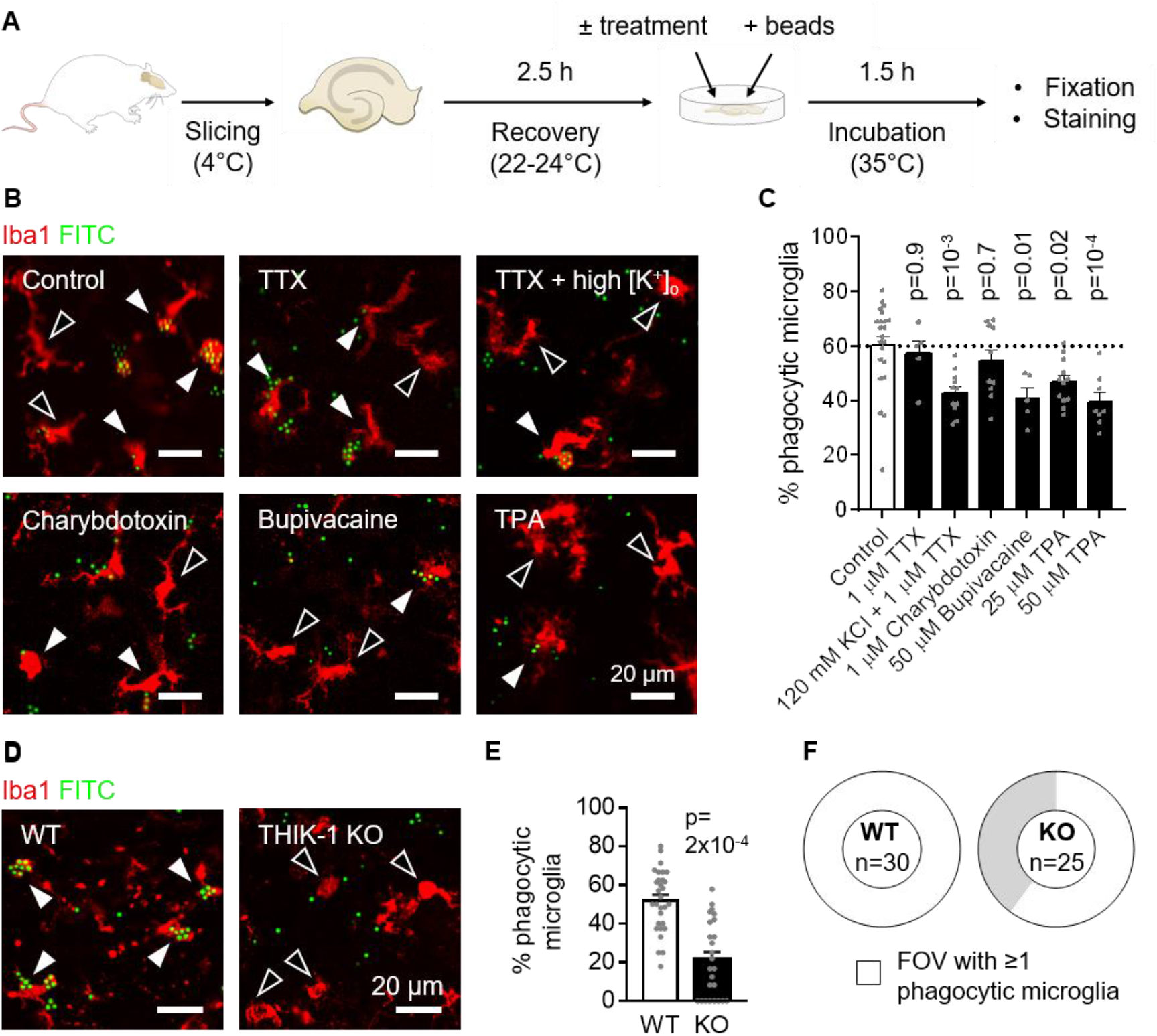
Two-pore domain K^+^ channels regulate microglial phagocytosis. (*A*) Diagram of the phagocytosis assay. (*B*) Representative single plane images of microglia (Iba1, red) in acute hippocampal rat slices incubated with 3 µm microbeads (FITC, green), either untreated (control) or treated with 1 µM tetrodotoxin (TTX), 1 µM TTX + 120 mM KCl (high [K^+^]_o_), 1 µM charybdotoxin, 50 µM bupivacaine or 50 µM tetrapentylammonium (TPA). Black arrowheads indicate non-phagocytic cells, white arrowheads indicate phagocytic cells. Some microbeads remain outside microglia. (*C*) Percentage of microglia that phagocytosed microbeads in each condition, showing a reduction by high [K^+^]_o_ and two-pore K^+^ channel block (bupivacaine, TPA) but not by calcium-activated K^+^ channel block (charybdotoxin) (control: n=22 slices from 6 animals; TTX: n=5 slices from 2 animals; TTX + 120 mM KCl: n=11 slices from 4 animals; charybdotoxin: n=10 slices from 4 animals; bupivacaine: n=5 slices from 2 animals; 25 µM TPA: n=12 slices from 3 animals; 50 µM TPA: n=8 slices from 3 animals). P-values compare with control and are corrected for multiple comparisons. (*D*) Representative single plane images of microglia in acute hippocampal slices from wildtype (WT) and THIK-1 knockout (KO) mice incubated with 3 µm microbeads. (*E*) Percentage of microglia that phagocytosed microbeads in each genotype, showing a reduction in the KO. (*F*) Doughnut charts showing that only 60% of the analysed fields of view (FOV) contained phagocytic microglia in the KO, while all did in the WT. The number of total microglia per FOV was not different (WT: 15.0 ± 1.8, KO: 12.3 ± 2.1; p=0.34) between genotypes.

**Fig. 2.**
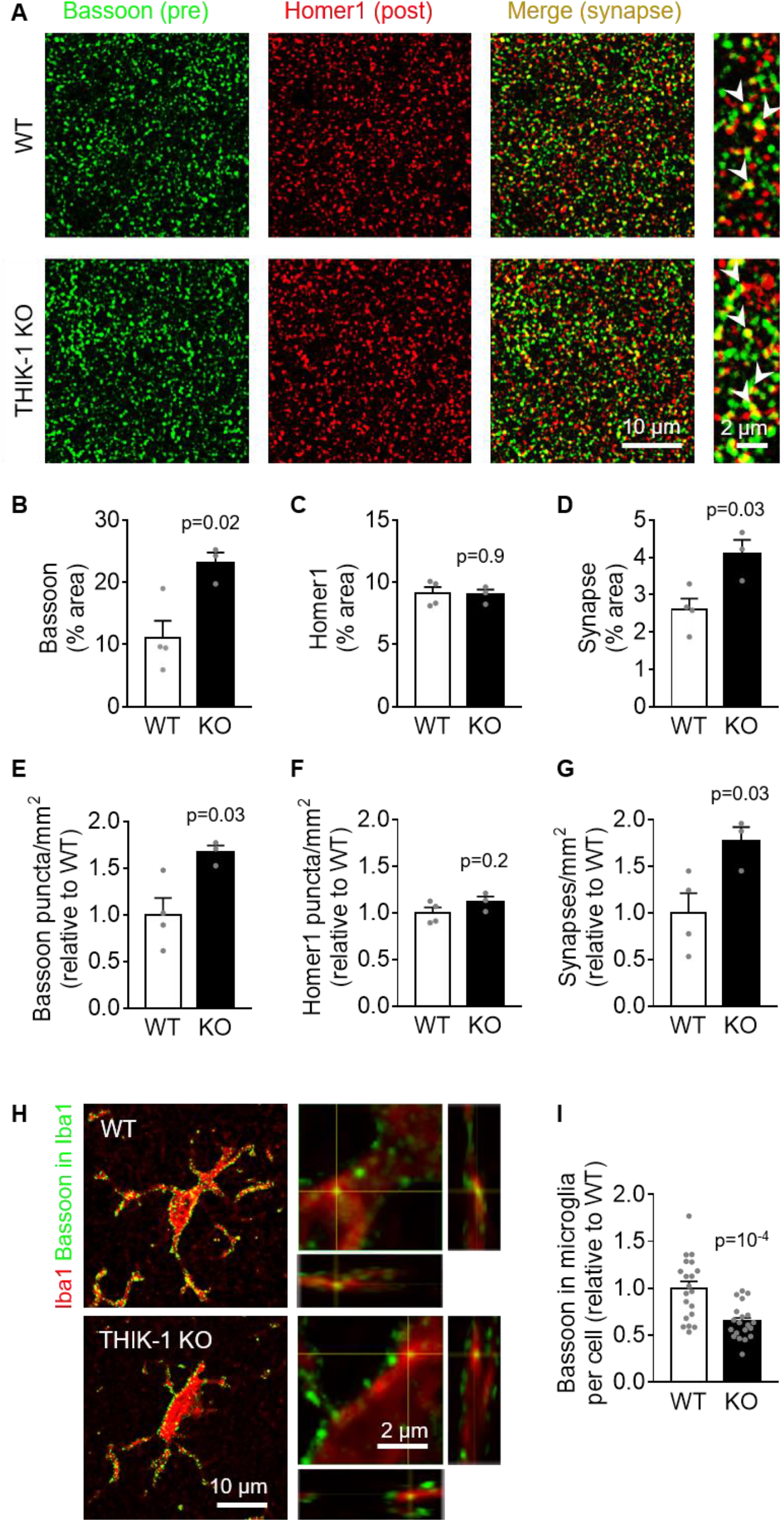
Microglial phagocytosis of presynaptic material is decreased in THIK-1 knockout mice. (*A*) Representative confocal images from the CA1 *stratum radiatum* of wildtype (WT) or THIK-1 knockout (KO) mice at P17-P19, showing the presynaptic marker Bassoon (green) and the excitatory postsynaptic marker Homer1 (red). The merged image and expanded views on the right show colocalized puncta (yellow). (*B-D*) Quantification of the area covered by (*B*) Bassoon, (*C*) Homer1 and (*D*) colocalized puncta, showing an increased fraction of the imaged area labeled for synapses in KO mice. (*E-G*) Quantification of the numbers of (*E*) Bassoon, (*F*) Homer and (*G*) synaptic puncta per mm^2^, showing increased numbers of presynaptic puncta and synapses in KO mice. Average numbers in WT mice were 53 puncta/100 µm^2^ for Bassoon, 42 puncta/100 µm^2^ for Homer1 and 26 synapses/100 µm^2^. (For panels B-G, WT: n=4 animals; KO: n=3 animals; 3 confocal stacks from 5 brain slices averaged per animal). (*E*) Representative confocal images showing Bassoon puncta (green) located within microglia (Iba1, red). On the right, close-up of microglial processes with orthogonal projections at the level of the crosshairs showing Bassoon puncta within microglia. (*F*) Quantification of the area of Bassoon puncta colocalizing with each microglial cell, showing a decrease in KO microglia (WT: n=20 cells from 4 animals; KO: n=20 cells from 3 animals).

## RESULTS

### Microglial phagocytosis is regulated by K^+^ channels

Ion channels and receptors controlling microglial motility might be involved in phagocytosis. These include P2Y_12_ receptors that regulate ATP-evoked microglial chemotaxis to an injury site (33) and THIK-1 K^+^ channels which regulate microglial morphology and surveillance (3). It was previously reported that P2Y_12_ receptors regulate phagocytosis (34, 35), but the role of microglial K^+^ channels is unknown. Therefore, we first tested whether blocking THIK-1 (the dominant K^+^ channel expressed in “resting” microglia) affects microglial phagocytosis *in situ* in brain slices. Following a recovery period of a few hours after brain slicing, which allows some microglial activation to occur (27), fluorophore-labeled resin microbeads were applied (28, 29) onto rat hippocampal slices in the presence or absence of pharmacological blockers (Fig. 1*A*).

Microglia engulf and phagocytose their substrates thanks to membrane protrusions and phagocytic cups, the formation of which relies upon cytoskeletal rearrangements (36). Indeed, microglia were seen to form phagocytic cups and engulf microbeads *in situ* (Fig. S1). As negative control experiments, the brain slices were incubated at 4°C, or in the presence of the actin inhibitor cytochalasin D, both of which disrupt cytoskeletal dynamics (37). We found that both conditions potently blocked phagocytosis (Fig. S2). MRS2578, a blocker of the P2Y_6_ receptor known to regulate microglial phagocytosis (38, 39) also reduced the fraction of microglia that were phagocytic (Fig. S2).

Potassium efflux across the microglial membrane via THIK-1 was previously found to control NLRP3 activation (3, 40). We found that elevating the extracellular K^+^ concentration ([K^+^]_o_) to prevent such efflux and depolarise the cells reduced the fraction of microglia that were phagocytic by a third (Fig. 1*B-C*). This was done in the presence of tetrodotoxin (TTX) to block voltage-gated Na^+^ channels and thus action potential-evoked synaptic transmitter release from neurons (TTX had no effect by itself: Fig. 1*C*). To determine how the raised [K^+^]_o_ altered phagocytosis, we applied blockers of different K^+^ channel types. Blocking Ca^2+^-activated K^+^ channels with charybdotoxin had no significant effect, but both the K^+^ channel blockers bupivacaine and tetrapentylammonium (TPA) inhibited phagocytosis (Fig. 1*B-C*). Since bupivacaine blocks both two-pore domain channels and voltage-gated Na^+^ channels while TPA blocks both two-pore domain channels and voltage-gated K^+^ channels (41, 42), these data are consistent with K^+^ efflux via THIK-1 (or downstream changes in microglial membrane voltage, V_m_) regulating phagocytosis. Since the pharmacology of these drugs is not entirely specific for THIK-1, we also used a mouse in which THIK-1 was knocked out (KO) and found that deletion of THIK-1 had a similar inhibitory effect on microglial phagocytosis (Fig. 1*D-F*).

### THIK-1 regulates microglial phagocytosis of synapses

Deficits in microglial phagocytosis could result in impaired pruning of synapses during development (43), so we tested the effects of THIK-1 KO on hippocampal synapse numbers. Using P17-P19 mice, when synapse pruning in the hippocampus is near its peak (12, 13), we assessed the labeling of presynaptic (Bassoon) and postsynaptic glutamatergic (Homer1) markers in the *stratum radiatum* of the CA1 hippocampal region by immunohistochemistry (30). A colocalization of both markers was taken to indicate an excitatory synapse (see Fig 2*A*). In the THIK-1 KO, while no postsynaptic change was detected, the fraction of the imaged area labeled by the presynaptic marker approximately doubled compared to that in wildtype littermates (WT). As a result, the derived total synaptic area was higher in KO mice (Fig. 2*B-D*). The increase in colocalization was produced by a 67% increase in the number (Fig. 2*E-G*), but not the size (Fig. S3), of presynaptic terminals, with no change in the number of postsynaptic terminals (Fig. 2*F*).

To demonstrate that the increase in presynaptic terminal number was caused by a reduction of phagocytosis, we next assessed the presence of Bassoon puncta inside Iba1-labeled microglia (Fig. 2*H*). In THIK-1 KO mice, Bassoon colocalization with microglia was significantly reduced (Fig. 2*I*). Taken together, these data suggest that THIK-1 regulates the number of glutamatergic synapses by promoting microglial uptake of presynaptic material.

### THIK-1 regulates excitatory synaptic transmission

To further confirm that THIK-1 regulates removal of functional excitatory synapses, we performed whole-cell voltage-clamp recording of CA1 pyramidal neurons from P17-19 mice (Fig. 3). We found that the spontaneous EPSC frequency was enhanced in pyramidal neurons from THIK-1 KO compared to WT mice, with no amplitude alteration (Fig. 3*A-C*). Since this increase in the KO could be due either to a higher number of excitatory synapses or to higher activity (or an increased vesicle release probability) of presynaptic neurons, we bath-applied TTX to block action potential-mediated neurotransmitter release. Consistent with our immunohistochemical studies showing more synapses in the KO (Fig. 2), mEPSC frequency approximately doubled in THIK-1 KO pyramidal neurons compared to in WT cells, without an amplitude change (Fig. 3*A*,*D-E*).

**Fig. 3.**
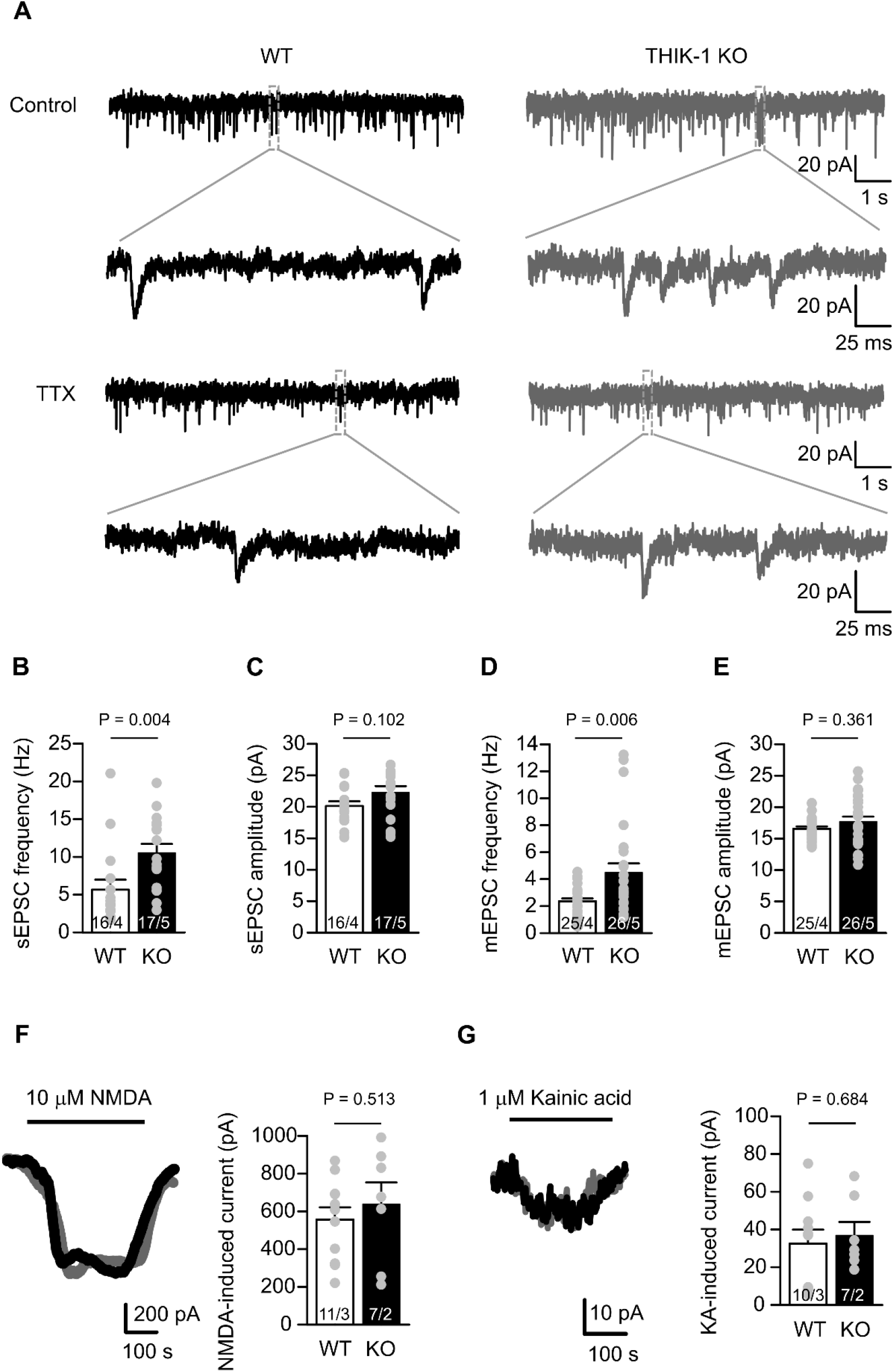
THIK-1 knockout increases excitatory synaptic transmission. (*A*) Representative current traces from whole-cell voltage-clamped CA1 pyramidal neurons (V_h_ = −65 mV, E_Cl_ = −62mV) in WT (black, left) and THIK-1 KO (grey, right) hippocampal slices. The top two rows show spontaneous EPSCs (sEPSCs) i.e., including both miniature EPSCs (mEPSCs) and those evoked by spontaneous action potentials. The bottom two rows show mEPSCs in 500 nM TTX. The second and fourth row traces expand the indicated areas in the first and third rows. (*B-C*) Bar graphs showing sEPSC (*B*) frequency and (*C*) amplitude, comparing WT and THIK-1 KO. Grey circles overlaid show raw values in individual cells. Numbers on bars are of cells/animals, and s.e.m. values use cells as the statistical unit. (*D-E*) As for C-D but showing mEPSC (*D*) frequency and (*E*) amplitude, comparing WT and THIK-1 KO in 500 nM TTX. (*F*) Representative traces of whole-cell currents (V_h_ = −40 mV, E_Cl_ = −62 mV) from WT (black) and THIK-1 KO (red) hippocampal CA1 pyramidal cells, and bar graph of mean current induced by 10 µM NMDA bath application to WT and THIK-1 KO cells. (*G*) As for (F) but applying 1 µM kainic acid.

This effect of THIK-1 deletion on synapse levels did not result from an altered number of microglia, as overall microglial density and distribution in CA1 were similar between THIK-1 WT and KO mice (Fig. S4). Furthermore, we found that THIK-1 only regulates excitatory synapses, with no detectable effect on inhibitory synapses. Neither the frequencies nor amplitudes of sIPSCs or mIPSCs were affected by THIK-1 KO (Fig. S5). Altogether, our data suggest that THIK-1 deficiency leads to an increase in the number of functional excitatory synapses, which is due to THIK-1 regulating microglial internalization of excitatory presynaptic terminals, presumably via its effect on the microglial membrane potential.

We tested postsynaptic effects of THIK-1 deficiency electrophysiologically (Fig. 3*F-G*) by bath-applying the glutamate receptor agonists NMDA (to activate NMDA receptors, 10 µM; Fig. 3*F*) and kainic acid (to activate AMPA/KA receptors, 1 µM; Fig. 3*G*). Consistent with our immunolabeling shwing no effect on postsynaptic terminals (Fig. 2*C*), we found no significant differences between the THIK-1 KO and WT in their NMDA- and kainate-induced macroscopic currents (Fig. 3*F-G*). Thus, THIK-1 deficiency selectively enhances the number of excitatory presynaptic release sites without affecting the postsynaptic glutamate receptor density (assessed from the spontaneous and miniature EPSC amplitudes) or the total (synaptic plus extrasynaptic) glutamate receptor density (assessed from the response to NMDA and kainate).

### Block of THIK-1 affects microglia and synapses in the human brain

Finally, to assess the relevance of our findings to understanding the human brain, we used live human cortical slices obtained from neurosurgically-resected biopsy tissue to study acute responses to THIK-1 blockers. Human microglia express THIK-1 at the RNA level (44, 45). Having confirmed that impairment of THIK-1 in rodents leads to microglial deramification (Fig. S6, in agreement with (3)) and shown that it increases the number of excitatory synapses (Fig. 3), we studied whether inhibiting THIK-1 pharmacologically (since KO experiments are not possible) could recapitulate these effects in live human microglia *in situ*. Acute (40-minute) block of THIK-1 channels with TPA significantly reduced microglial process length and ramification, as revealed by three-dimensional Sholl analysis (Fig. 4*A-D*). Perhaps surprisingly, given the brief period of block, the fractional area of the image that was labeled for the vesicular glutamate transporter 1 (vGluT1), an excitatory presynaptic marker, was increased by 25% in the TPA-treated slices (Fig. 4*E-F*). We obtained similar effects on microglial ramification and synapse number when treating slices with bupivacaine (Fig. S7). Bupivacaine and TPA have different targets but both block two-pore K^+^ channels (see above), consistent with THIK-1 mediating the effects seen (although we cannot rule out the possibility that these drugs share another, unreported target). Taken together, these data suggest that THIK-1 channels control microglial morphology and synapse levels in the adult human brain.

**Fig. 4.**
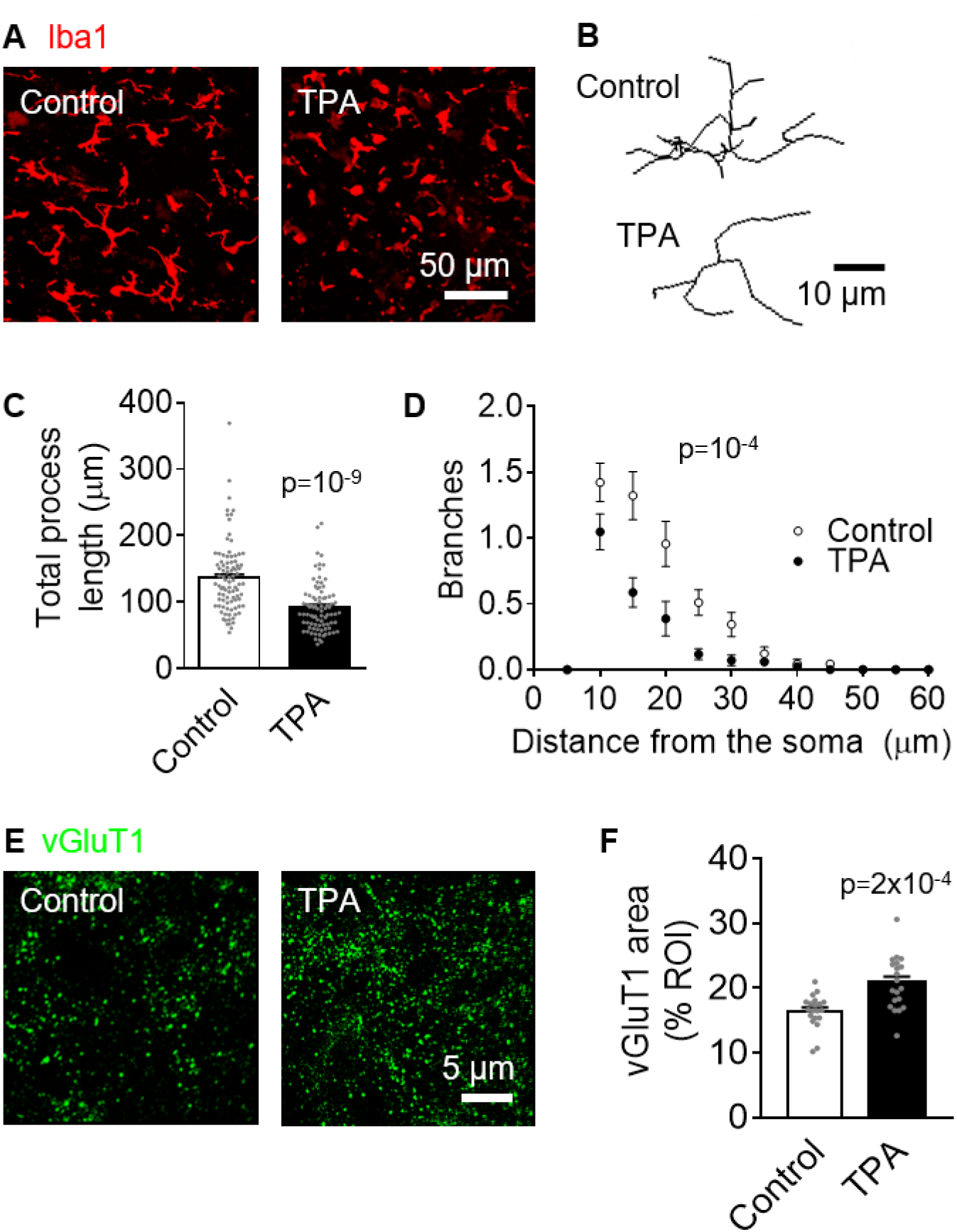
Block of THIK-1 deramifies microglia and increases presynaptic area in human brain slices. (*A*) Representative confocal images from human cerebral cortical slices showing the effect of 50 µM TPA treatment (40 mins) on microglial morphology (Iba1, red), which decreased microglial ramification and process length. (*B-D*) Morphological analysis of microglia from control (90 cells from 3 human subjects) and TPA-treated microglia (85 cells from 3 human subjects), showing (*B*) representative 3D-reconstructed microglia, and Sholl analysis-derived (*C*) total process length and (*D*) number of process branches at distances (in 5 μm increments) from the cell soma. (*E*) Representative SRRF-confocal images showing vGluT1 puncta (green). (*F*) Quantification of the area covered by vGluT1, showing an increase in TPA-treated slices compared to controls (20 slices from 2 human subjects per condition).

## DISCUSSION

Microglia are not merely passive support cells. Instead, they continuously survey the brain parenchyma and control neuronal function. THIK-1 channels, the main K^+^ channels in “resting” microglia, regulate their ramification, surveillance and cytokine release (3). Here, using pharmacology and THIK-1 knock-out mice, we demonstrated that microglial phagocytosis is also controlled by THIK-1 (Fig. 1). As a result, both immunohistochemistry and electrophysiology show that the number of hippocampal excitatory synapses is increased in THIK-1 KO mice (Fig. 2, Fig. 3).

The requirement of THIK-1 for phagocytosis may in part be due to its role in enhancing microglial ramification and surveillance, which will increase the probability of a microglial cell encountering a target to phagocytose. However, the tonically active THIK-1 may also promote phagocytosis by keeping microglia hyperpolarized (3), since a hyperpolarized membrane voltage is associated with phagocytosis in macrophages (20). A depolarized membrane potential is also seen in TREM2 KO microglia, in which phagocytosis is reduced (21).

Our study indicates that THIK-1-mediated promotion of synapse loss is mainly via a presynaptic effect (Fig. 2*B-D*). While others have reported internalization of postsynaptic materials as well (13, 46), there is now evidence that microglial phagocytosis preferentially targets presynaptic compartments both in health (12, 14) and disease (25, 26, 47). This might be partly due to “eat-me” tags (such as C1q (48)) being preferentially located on presynaptic sites. On the other hand, our data indicate that deleting THIK-1 has no effect on inhibitory synapses (Fig. S5), suggesting that phagocytosis by microglia mainly targets excitatory synapses, and thus that there is a difference in the recognition molecules expressed on excitatory versus inhibitory synapses. Indeed, microglial depletion increases mEPSC frequency in brain slices (49) and *in vivo* (50), while mIPSCs remain unaltered (50).

Adult rodent (51) and human (52) microglia continue to internalize synapses into adulthood, after the normal developmental period is over, which might be detrimental if excessive (53). In mice, memories persist better when microglia are depleted (51). Adult human microglia express *Kcnk13*, the gene encoding THIK-1 (44, 45) and we found that even short-term application of THIK-1 blockers increased presynaptic vGluT1 in live human brain slices (Fig. 4, Fig. S7), which suggests a fast removal (and presumably replacement) of synapses. Work on hippocampal slices has shown that individual presynaptic internalization events are rapid (frequently shorter than 3 minutes (12)). However, turnover of synapses is generally low *in vivo*, taking from hours to months (54–56), which contrasts with our data (Fig. 4*E-F*) that suggest robust replacement in acute brain slices. Synapse turnover may be much faster *ex-vivo* due to microglia being partly activated and neurons being damaged by the slicing procedure (57). Indeed, slicing leads to loss of spines and synaptic responses (58). This parallels the acceleration of synapse loss reported in traumatic brain injury in humans, although that occurs over days (59). Despite these differences with the *in vivo* situation, our results suggest a role of microglial THIK-1 in regulating human synapse turnover.

Synapse loss is a strong indicator of cognitive decline (60–62). Synaptic deficits precede amyloid deposition in animal models of dementia (63) and microglial phagocytosis of synaptic material is increased in Alzheimer’s patients (52). Ablation of microglia rescues synaptic loss and reduces memory impairment in mouse models of dementia (64). Thus, manipulating microglia-synapse interactions may provide clinical benefit for cognitive impairment. Specifically, being able to block microglial phagocytosis in a time-controlled manner could help protect synapses from removal. It would be interesting to test whether short-term block of THIK-1 *in vivo* (e.g., using THIK-1-blocking anaesthetics such as isoflurane or sevoflurane) results in increased synapse number, and the magnitude and duration of any such effect in normal ageing, brain injury and dementia models. Microglial responses (and especially phagocytosis) are crucial for the development and progression of dementia (65). Since amyloid-targeting therapies have all failed, possibly because therapeutic interventions are given too late (60), it would be advantageous to devise therapeutic agents that control phagocytosis to prevent synapse loss early on.

## ACKNOWLEDGEMENTS

This work was supported by European Research Council (BrainEnergy) and Wellcome Investigator Awards (099222) to DA, a Wellcome Trust four-year PhD studentship to PI and a Chulabhorn Royal Academy PhD studentship to CH. For the purpose of Open Access, the authors have applied a CC-BY public licence to any Author Accepted Manuscript version arising from this submission. We thank Anna Barkaway, Damian Cummings, Frances Edwards, Soyon Hong, Vasiliki Kyrargyri, Jonathan Lezmy, Yuening Li, Christian Madry, Thomas Pfeiffer, Tania Quintela-López, Patricia Salinas and James Scott-Solache for comments on the manuscript.

## DATA AVAILABILITY

All data and code are available upon reasonable request.

## Supplementary Information

**Fig. S1.**
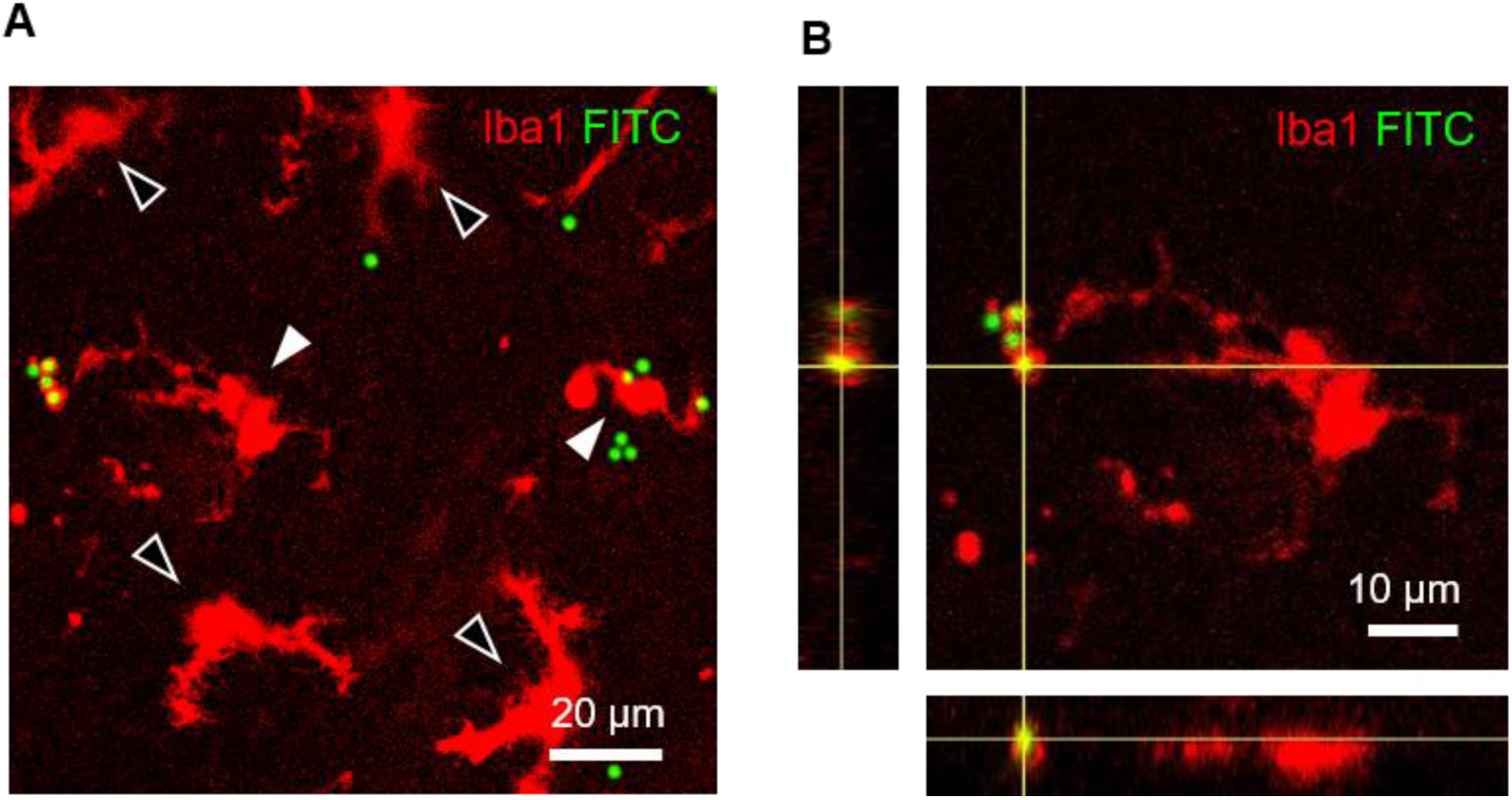
Microglia phagocytose fluorophore-labeled microbeads. (*A*) Representative maximum intensity projection of a confocal z-stack of microglia (Iba1, red) in acute hippocampal rat slices incubated with 3 µm microbeads (FITC, green). Black arrowheads indicate non-phagocytic cells, white arrowheads indicate phagocytic cells. Some microbeads remain outside microglia. (*B*) Close-up of the phagocytic microglial cell at the left of panel A. Orthogonal projections at the level of the crosshairs show engulfment of the fluorescent microbeads inside a phagocytic cup.

**Fig. S2.**
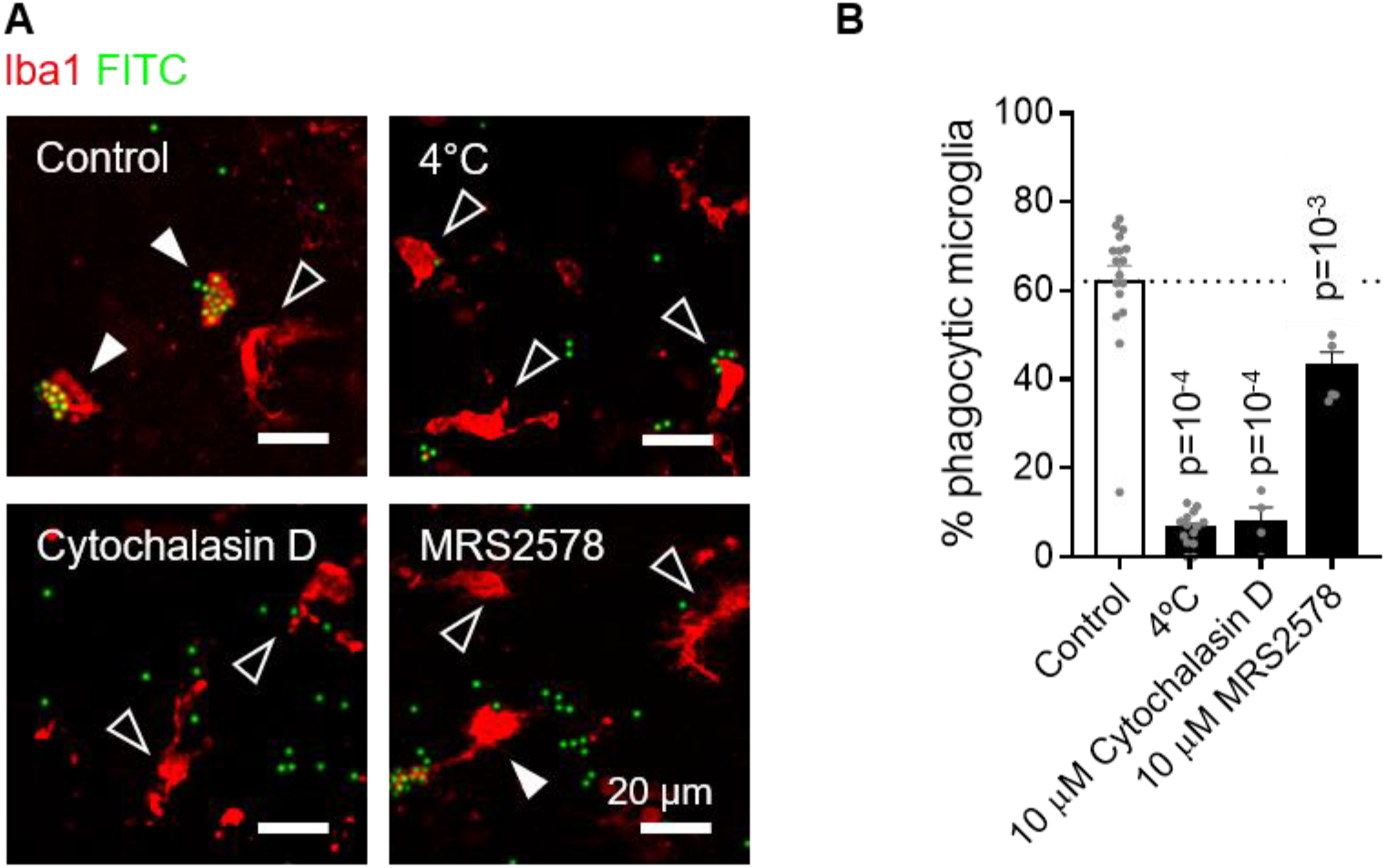
Pharmacological inhibition of microglial phagocytosis. (*A*) Representative single plane images of microglia (Iba1, red) in acute hippocampal rat slices incubated with 3 µm microbeads (FITC, green), at 35°C (control, 10 µM cytochalasin D or 10 µM MRS2578 (a P2Y_6_ blocker)) or 4°C. Black arrowheads indicate non-phagocytic cells, white arrowheads indicate phagocytic cells. (*B*) Percentage of microglia that phagocytosed microbeads in each condition, showing a reduction by exposure to cold (4°C), cytoskeletal disruption (cytochalasin D) or block of P2Y_6_ receptors (MRS2578) (control: n=17 slices from 4 animals; 4°C: n=14 slices from 4 animals; cytochalasin D: n=4 slices from one animal; MRS2578: n=6 slices from 3 animals).

**Fig. S3.**
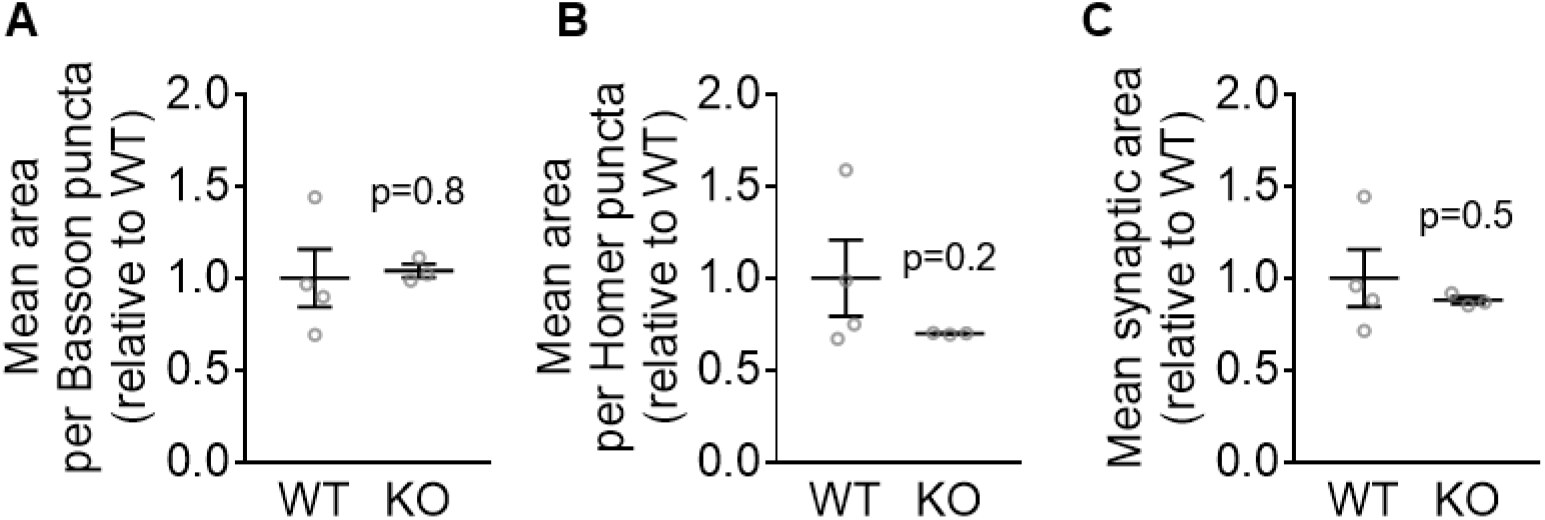
Synaptic terminal size is not affected in THIK-1 knockout mice. (*A-C*) Quantification of (*A*) Bassoon, (*B*) Homer and (*C*) overlapping puncta size, showing no changes in KO mice (WT: n=4 animals; KO: n=3 animals; 3 confocal stacks from 5 brain slices averaged per animal).

**Fig. S4.**
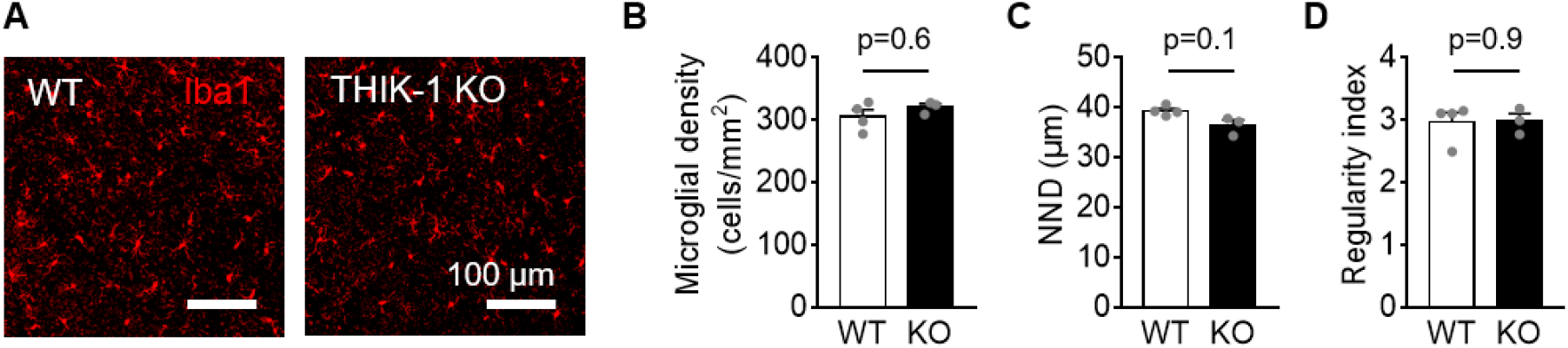
Microglial density and spatial distribution are not changed in CA1 area of THIK-1 knockout mice. (*A*) Representative maximum intensity projection of confocal z-stacks showing hippocampal microglia (Iba1, red) in wildtype (WT) and THIK-1 knockout (KO) mice. (*B-D*) Quantification of (*B*) microglial density, (*C*) average nearest-neighbour distance (NND) and (*D*) regularity index, showing no differences between genotypes (WT: n=4 animals; KO: n=3 animals; 5 slices averaged per animal).

**Fig. S5.**
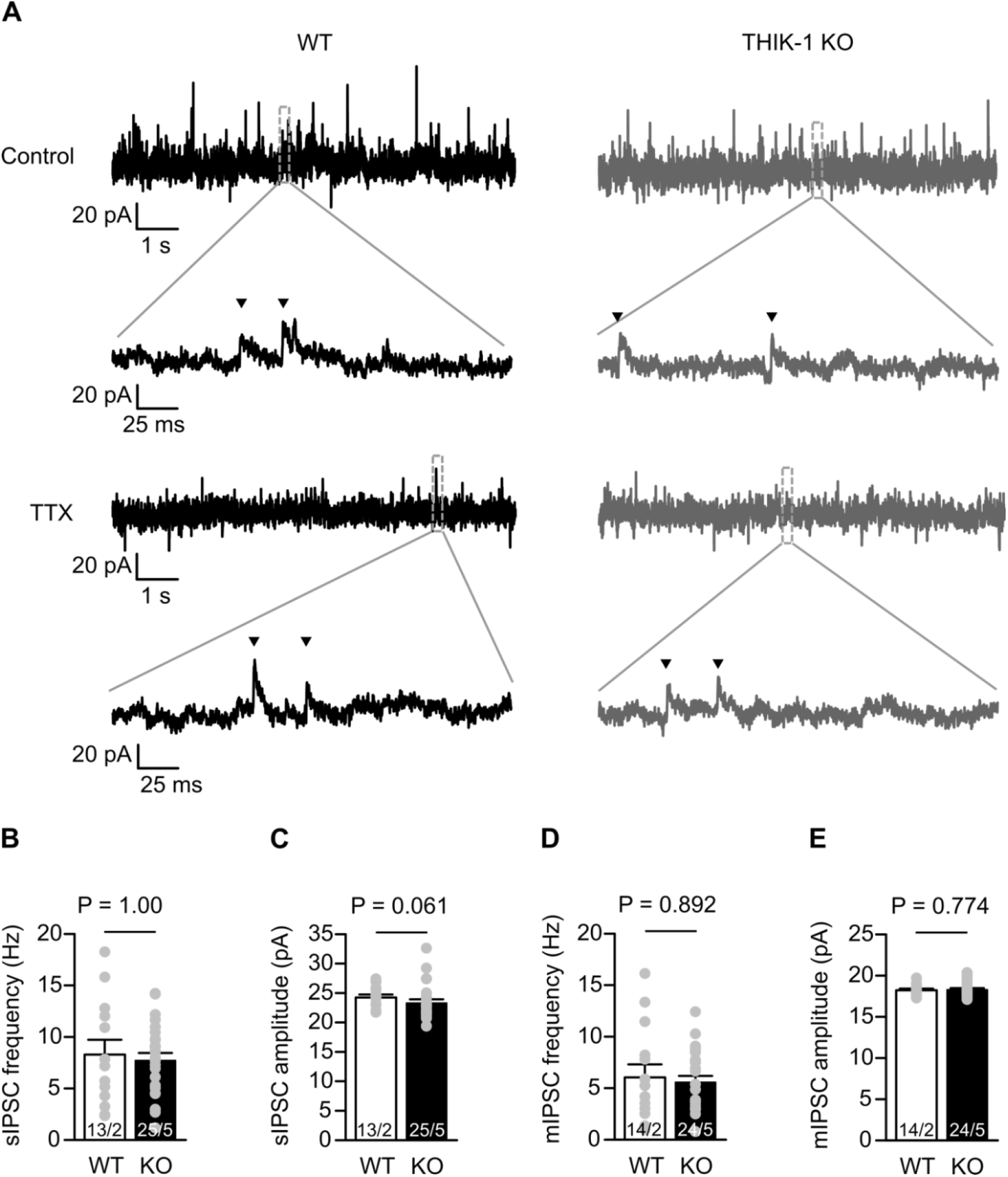
THIK-1 knockout does not alter inhibitory synaptic transmission. (*A*) Representative current traces from whole-cell voltage-clamped CA1 pyramidal neurons (V_h_ = −50 mV, E_Cl_ = −85mV) in WT (black, left) and THIK-1 KO (grey, right) hippocampal slices obtained from P26–32 mice. Top two row traces show spontaneous IPSCs (sIPSCs, highlighted with triangles), and bottom two row traces show miniature IPSCs (mIPSCs) in 500 nM TTX. The second and fourth row traces expand the boxed areas in the first and the third rows. (*B-C*) Bar graphs showing sIPSC (*B*) frequency and (*C*) amplitude, comparing WT and THIK-1 KO. Grey circles overlaid show raw data. Numbers on bars are of cells/(animals), and s.e.m. values use cells as the statistical unit. (*D-E*) As for C-D but showing mIPSC (*D*) frequency and (*E*) amplitude, comparing WT and THIK-1 KO in 500 nM TTX.

**Fig. S6.**
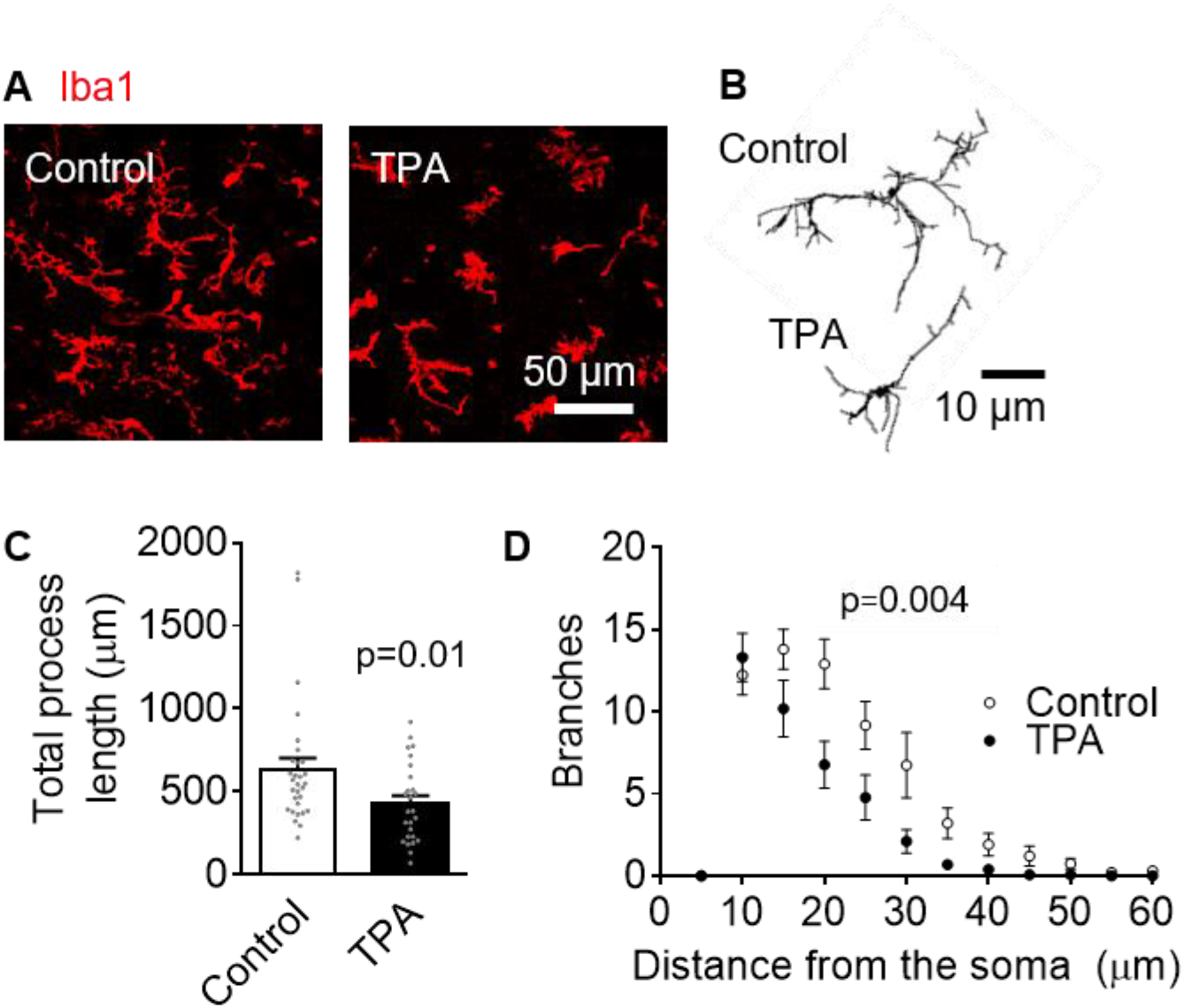
Block of THIK-1 deramifies microglia in rat brain slices. (*A*) Representative confocal images from rat hippocampal slices showing the effect of 50 µM TPA treatment (40 mins) on microglial morphology (Iba1, red), which decreased microglial ramification and process length. (*B-D*) Morphological analysis of microglia from control (30 cells from 2 animals) and TPA-treated microglia (26 cells from 2 animals), showing (*B*) representative 3D-reconstructed microglia, and Sholl analysis-derived (*C*) total process length and (*D*) number of process branches at distances (in 5 μm increments) from the cell soma.

**Fig. S7.**
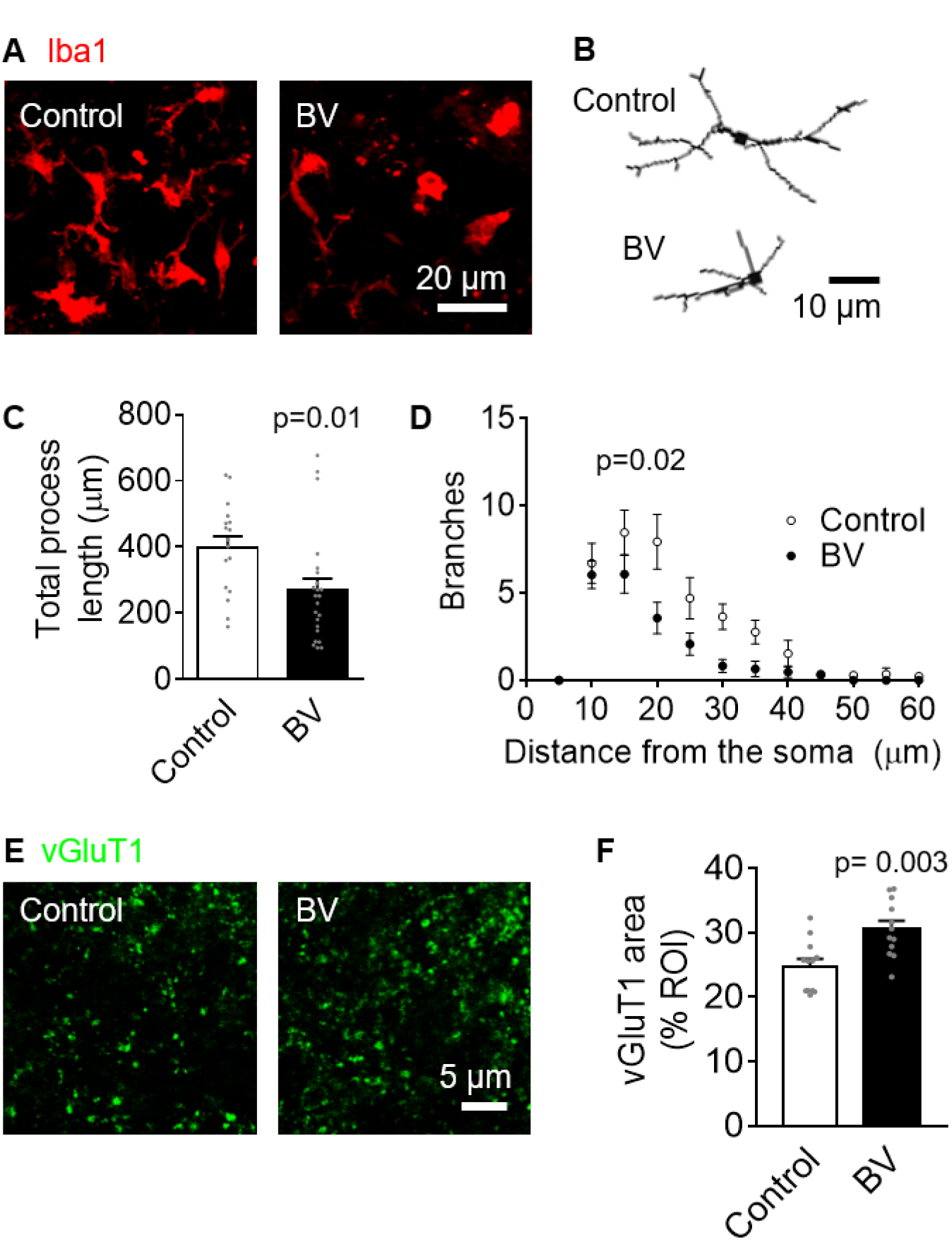
Block of THIK-1 with bupivacaine deramifies microglia and increases presynaptic area in human brain slices. (*A*) Representative confocal images from human cerebral cortical slices showing the effect of 50 µM bupivacaine treatment (40 mins) on microglial morphology (Iba1, red), which decreased microglial ramification and process length. (*B-D*) Morphological analysis of microglia from control (17 cells from one human subject) and bupivacaine-treated microglia (23 cells from one human subject), showing (*B*) representative 3D-reconstructed microglia, and Sholl analysis-derived (*C*) total process length and (D) number of process branches at distances (in 5 μm increments) from the cell soma. (E) Representative SRRF-confocal images showing vGluT1 puncta (green). (F) Quantification of the area covered by vGluT1, showing an increase in bupivacaine-treated slices compared to controls (12 per condition from one human subject).

